# A structure-guided antibody detects SOD1 oligomers in diverse ALS genotypes

**DOI:** 10.1101/2025.05.05.652290

**Authors:** Smriti Sangwan, Hannah E. Rieder, Destaye Moore, Negar Khanlou, Bennett Novitch, Mark Geisberg, David S. Eisenberg

## Abstract

Antibodies offer versatility as diagnostic and therapeutic tools to target specific protein epitopes. However, the transient nature of intermediate protein conformations, such as that of amyloid oligomers, poses a challenge for antibody development. We use a structure-guided approach to generate a monoclonal antibody against oligomers of Superoxide Dismutase 1 (SOD1). Mutations in SOD1 are linked to a subset of familial Amyotrophic Lateral Sclerosis (fALS), a fatal neurodegenerative disease. Based on the corkscrew-like features of non-native SOD1 oligomers previously determined, we generate an antibody specific to SOD1 oligomers. We show that the antibody, CSAb detects SOD1 oligomers, not fibrils or native SOD1, and alleviates the cytotoxic effects of SOD1 oligomers in a cell culture model of primary motor neurons. Immunohistochemical analyses of human ALS subjects show CSAb reactivity in both neuronal and non-neuronal cells. Finally, we provide evidence that CSAb reactive SOD1 oligomers are present in non-SOD1 linked fALS and sporadic ALS subjects. Together, our study provides a new probe against SOD1 oligomers and suggests that cytotoxic SOD1 oligomers are prevalent in diverse ALS genotypes.

## Introduction

The conversion of soluble proteins into insoluble amyloid fibrils underlies the pathogenicity of neurodegenerative diseases such Alzheimer’s, Parkinson’s, and Amyotrophic Lateral Sclerosis (ALS). In each disease, proteins convert from their native structure to an ensemble of non-native conformations ranging from small oligomers to insoluble amyloid fibrils. Several lines of evidence suggest that amyloid oligomers possess cytotoxic properties, while fibrils may be agents of pathogenic propagation through seeding^1–4^. Current therapeutic strategies, including gene and immune therapies, have largely been unsuccessful, and there is a need for tools that detect and target oligomer and fibril conformations in neurodegenerative diseases.

Antibodies are among the most promising approaches for treatment and characterization of neurodegenerative diseases. The functional versatility of antibodies enables their use as therapeutics by blocking the target epitope and as probes to detect distinct conformational epitopes. The ability to increase their efficacy with maturation methods allows for the development of ultra-high affinities as has been shown for cell receptors and infectious proteins^5,6^. However, developing antibodies against amyloid conformations has been challenging. Traditionally, antibodies have been developed by inoculating animals with proteins and screening for desired candidates from the sera. This approach is not suitable for selecting amyloid intermediates, such as oligomers, due to their transient nature. Indeed, known amyloid antibodies provide modest therapeutic benefit in animal models, and their precise epitopes are unknown, limiting efforts to improve their binding^7–9^. Recently, we utilized a structure-guided approach to develop nanobodies^10^. The nanobodies were grafted with peptide-based inhibitors in their complementarity-determining regions. We showed that the nanobodies bind fibrils of Alzheimer’s associated protein, Tau, and prevent its seeded propagation. However, this method cannot be used to develop antibodies against amyloid oligomers due to the limited structural information available on them. An alternative strategy is to use shorter segments as immunogens. These shorter amyloid segments often adopt stable conformations with well-defined structural features. This allows for the development of antibodies that recognize conformational epitopes rather than the amino acid sequence.

Superoxide dismutase 1 (SOD1) is a potential candidate for therapeutic development in ALS. Wild-type SOD1 (SOD1^WT^) is a metal-binding homodimer that protects cells against oxidative damage^11^. ALS-associated mutations destabilize the protein, rendering it prone to misfolding and aggregation. Therapeutic strategies targeting SOD1-linked ALS are currently being explored, including antisense oligonucleotides (ASOs) that suppress SOD1 expression and small molecules designed to stabilize the native conformation of SOD1 and prevent misfolding ^12,13^. These approaches have shown encouraging results in clinical trials, suggesting that efforts to block SOD1 aggregation are a viable therapeutic strategy.

While the role of SOD1 in fALS is clear, several studies implicate SOD1 in other ALS forms. SOD1 aggregates have been detected through immunohistochemistry (IHC) in spinal cord tissues from sALS and non-SOD1 fALS patients as well as through biochemical analysis of patient-derived lymphoblasts^14–16^. Over-expression of SOD1^WT^ in mice leads to neuronal loss and exacerbates disease progression^17^. However, whether WT and familial mutants share the same aggregation pathway and cause neuronal dysfunction through a common mechanism is unclear. Cryo-electron microscopy (cryoEM) studies of SOD1^WT^ fibrils and familial mutants show different fibril architectures, suggesting potential differences in their aggregation pathway that may differentiate SOD1 aggregates in different ALS forms^18,19^. Together, these findings highlight a potentially broad role of SOD1 in ALS disease pathogenesis and the need to develop novel approaches to detect and prevent the formation of cytotoxic conformations of SOD1.

Current antibodies used to detect non-native SOD1 conformations in ALS lack specificity. Antibodies such as C4F6 and B8H10 cannot distinguish between fibrillar and oligomeric forms, and their epitope recognition is poorly characterized, leading to inconsistent results in detecting toxic species^20,21^. The inability to precisely target specific toxic conformations of SOD1 limits our understanding of ALS pathogenesis and the development of reliable biomarkers for diagnosis and disease progression. Furthermore, it limits the potential of immunotherapy strategies aimed at selectively neutralizing toxic SOD1 conformations while preserving its normal function.

The overlap of SOD1 in both fALS and sALS has prompted growing interest in the specific structural and conformational changes in SOD1 that may underlie its pathogenicity in ALS. Recent studies suggest that it is not the large fibrillar aggregates of mutant SOD1 but rather small, soluble oligomers that may be the primary toxic species in ALS ^22–25^. The oligomeric forms of SOD1 are thought to adopt non-native conformations that expose buried hydrophobic surfaces, leading to abnormal interactions with cellular components and disrupting cellular homeostasis. In our previous work, we uncovered a unique structural element within SOD1 that appears critical for its oligomeric toxicity. Specifically, we identified residues 28-38 of SOD1 as forming a “corkscrew” motif—a twisted β-sheet comprised of antiparallel out-of-register β-strands—normally buried within the native SOD1 structure^24^. We proposed that when SOD1 is destabilized, such as through mutations or cellular stressors, the corkscrew segment becomes exposed, enabling oligomerization^24,26^. The formation of these oligomers is associated with neurotoxicity, as they disrupt cellular homeostasis and contribute to motor neuron degeneration.

## Results

### Design of a corkscrew-specific antibody

We hypothesized that the corkscrew segment could be used to develop an antibody specific for SOD1 oligomers. Based on the corkscrew structure, we proposed a model whereby residues Val29, Val31, Gly33, and Ile35, which remain buried in native folded SOD1, become solvent-exposed upon misfolding and promote self-assembly into oligomers (Fig. 1A, 1B). We hypothesized that we could utilize this difference in solvent accessibility to develop an antibody specific for the corkscrew conformation. To generate the antibody, mice were immunized with the synthetic peptide KVKVWGSIKGL corresponding to the SOD1 (28-38) epitope with a P28K substitution previously used to increase the solubility of the segment. Following immunization and booster injections, hybridoma technology was used to create monoclonal antibody-producing cells. Screening was performed via ELISA and dot blots to isolate a monoclonal antibody hereby named CSAb (Corkscrew antibody). The final antibody was purified from bulk culture using Protein G chromatography.

**Figure 1:**
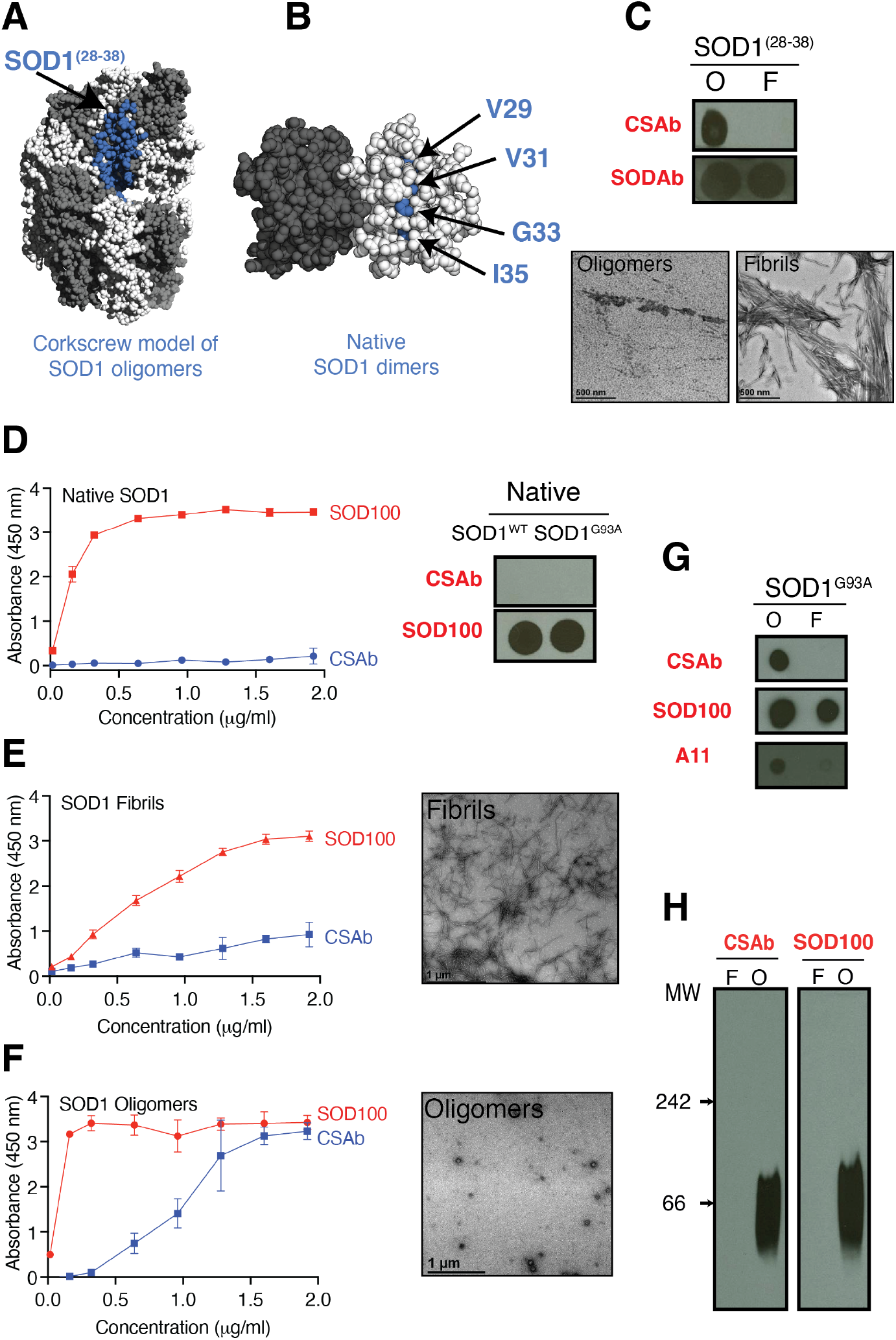
An oligomer-specific SOD1 antibody. (A) Model of full-length SOD1 oligomers based on the corkscrew structure adopted by segment 28-38. Notice that segment 28-38 (colored blue) is solvent exposed.**(B)** Structure of wild-type SOD1 as a dimer (PDB:2C9S). Residues Val29, Val31, Gly33, Ile35 of segment 28-38 (colored blue) are buried in the core. **(C)** Dot blot assay showing CSAb recognizes oligomeric forms of segment 28-38 but not the fibrillar form. SOD100 was used as loading control and shows equivalent binding to both forms of the segment. Oligomers and fibrils were generated by aggregating the segment for 4 and 12 hours, respectively. Negative stain micrographs of the oligomeric and fibrillar preparations of the segment show abundant fibrils (bottom). **(C)** ELISA shows CSAb does not bind native metal-bound SOD1^WT^ protein. SOD100 was used as a control. Dot blot assay with both WT and G93A mutant protein showing CSAb does not bind native proteins. **(E**,**F)** ELISA showing CSAb binds to SOD1 oligomers. Negative stain micrographs confirm the presence of abundant fibrils in the fibrillar sample and smaller oligomers in the oligomer sample. **(G)** Dot blot assay showing CSAb binds to SOD1G93A oligomers that are recognized by the amyloid oligomer antibody, A11. SOD100 was used as control. **(H)** Native gel electrophoresis followed by immunolotting with CSAb show oligomers range from 40-100 Kda, corresponding to dimers and hexamers.

### CSAb recognizes a conformational epitope presented by SOD1^28-38^

To investigate the specificity of CSAb, we conducted a series of assays comparing its reactivity to different assemblies of the segment in isolation and the full-length protein. To this end, we prepared oligomeric and fibrillar forms of the segment SOD1^28-38^ and measured CSAb’s reactivity using dot blot assays. CSAb detected oligomers but not fibrils of SOD1^28-38^ (Fig. 1C). Next, we tested CSAb against different conformational states of full-length SOD1. The native metal-bound protein was expressed and purified recombinantly, while fibrillar and oligomeric forms were generated using previously established protocols involving Cu^2+^ and Zn^2+^ ion removal and agitation-induced aggregation^24,27^. CSAb recognizes the oligomeric forms of apo-SOD1 but not the native metal-bound protein or the fibrillar forms (Fig. 1D-1F). Next, we compared CSAb’s reactivity to A11, a polyclonal antibody that detects oligomers of amyloid proteins regardless of sequence^28^. Similar to A11, CSAb detected SOD1 oligomers but not SOD1 fibrils (Fig. 1F). Blue-native gel electrophoresis followed by western blotting demonstrated that CSAb binds to soluble oligomers, which range in size from approximately 30 to 100 kDa, or about 2-6 monomers (Fig. 1G). CSAb also does not recognize the G33W mutant SOD1, a variant that disrupts the corkscrew conformation (Fig S1C). Collectively, these results suggest that CSAb recognizes a conformational epitope exposed specifically in the oligomeric forms of SOD1.

To further define the epitope recognized by CSAb, we performed immunoblotting under denaturing conditions. We hypothesized that under denaturing conditions, the protein’s secondary and tertiary structure would be disrupted and immunobinding assays would reveal if CSAb recognizes a specific conformation or the amino acid sequence of SOD1^28-38^. Western blotting after denaturing SDS-PAGE revealed that CSAb does not bind metal-bound SOD1 recombinantly purified or extracted from cell lysates (Fig. S1A). Interestingly, CSAb recognized apo-SOD1 in denatured conditions, indicating that CSAb’s binding under denaturing conditions may be influenced by metal availability (Fig. S1B). Taken together, these results suggest that CSAb recognizes a conformational epitope presented by the segment 28-38 in its apo-form.

### CSAb neutralizes the toxicity of SOD1 oligomers

Given that CSAb recognizes SOD1 oligomers, we hypothesized that it should be able to neutralize their cytotoxicity. We tested our hypothesis using a cell culture assay of primary motor neurons labeled with eGFP (enhanced green fluorescent protein) that we had previously used to show that corkscrew oligomers are cytotoxic ^24^. In this assay, SOD1 oligomers applied exogenously to motor neurons cause visible loss of dendrites and eventual loss of cell viability. We prepared full-length SOD1 oligomers and incubated them with sub-stoichiometric amounts of CSAb and applied them to cells in culture. Morphological analysis of motor neurons revealed that cultures treated with CSAb maintained neuronal integrity and healthy cell bodies, while those treated with the control IgG1 or buffer exhibited signs of toxicity, including shrinkage and loss of processes (Fig. 2A). Cellular viability was also partially rescued as measured by the MTT cell viability assay (Fig. 2B). To further investigate the cellular distribution of SOD1 oligomers and their potential mechanisms of toxicity, we visualized them using immunofluorescent staining with CSAb. Immunostaining revealed bright, punctate deposits most prominently localized near motor neuron cell bodies, where they appeared larger and more concentrated (Fig. 2C). A more diffuse pattern of SOD1 staining was observed in surrounding glial cells, suggesting that SOD1 may exert its toxic effects through the disruption of glial cell function. These findings demonstrate the importance of the 28-38 segment as a critical determinant of cytotoxicity and highlight how targeting this region with CSAb could mitigate its toxic effects.

**Figure 2:**
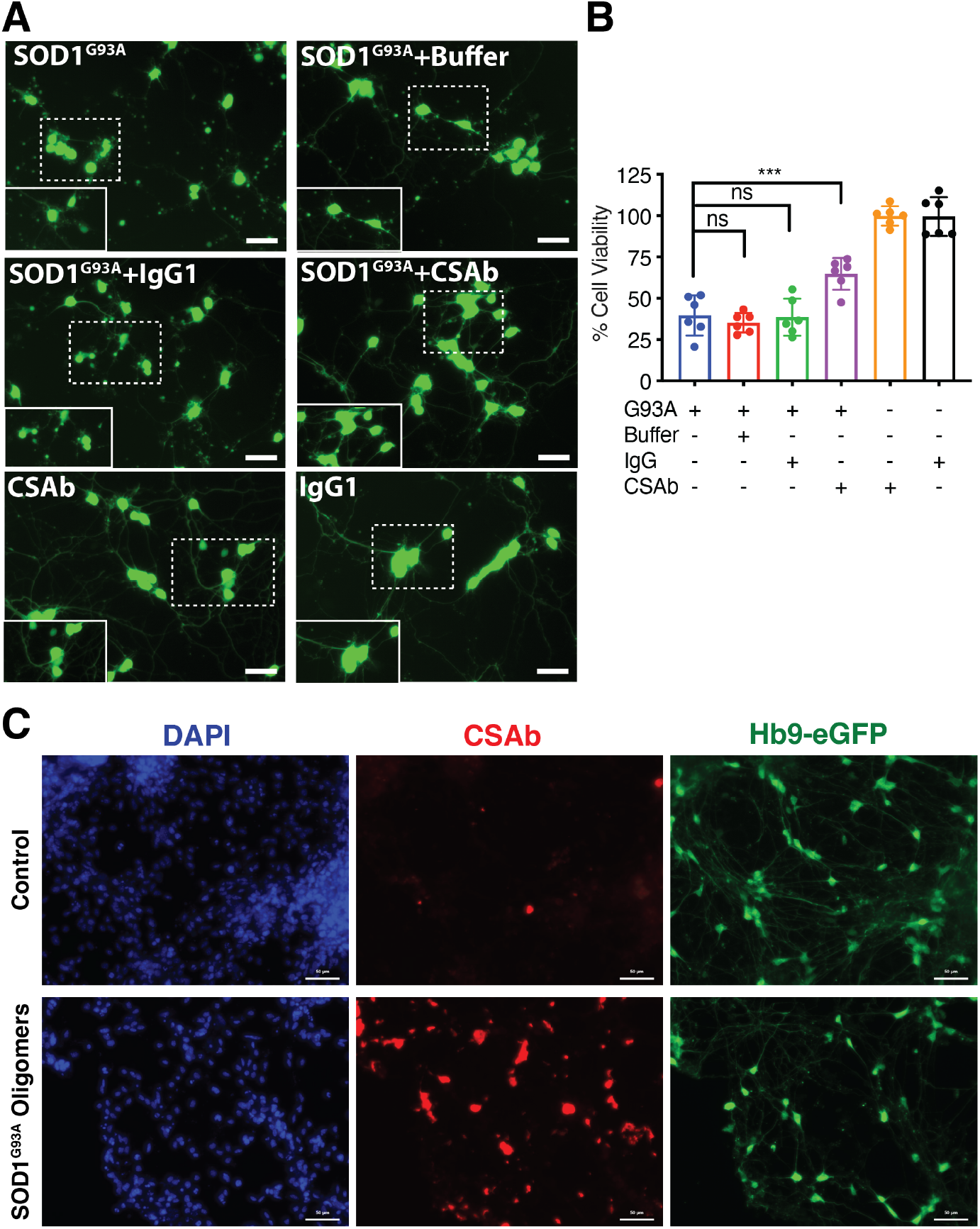
Co-incubation of SOD1 G93A oligomers with CSAb reduces their cytotoxic effects on cultured motor neurons. **(A)** Cultured motor neurons expressing eGFP were treated with 8 µM SOD1^G93A^ oligomers. Oligomers were incubated with CSAb and an unrelated IgG1 before applying to motor neurons. Cells treated with SOD1^G93A^ oligomers (top row left) displayed loss of neurites and rounding of cell bodies. Treatment with CSAb (middle row, right) alleviated the toxic effects and cells displayed normal physiology. **(B)** Cell viability measured by MTT assay showed partial rescue of viability by CSAb compared to buffer or an unrelated IgG. Results shown as mean and standard deviation. Statistical significance was determined by ANOVA. ns not significant, *** <.001. **(C)** Motor neurons treated with SOD1 G93A oligomers were immunostained with CSAb. Large oligomers were immunostained but not the endogenous folded SOD1WT.

### CSAb detects non-native oligomers in ALS patient tissues

Having determined that CSAb detects cytotoxic SOD1 oligomers, we asked if these oligomers are present in human ALS subjects. We performed immunohistochemistry on ALS patient tissues and compared them to healthy controls (clinical information in supplementary table 1). CSAb positive staining was observed in the motor neurons of four out of six SOD1-linked fALS cases, three out of six sALS, and four out of five non-SOD1 fALS (Fig. 3A, 3B, S2). We confirmed the specificity by comparing ten healthy human subjects and observed no detectable staining in any healthy subjects. To ensure that the observed reactivity was not due to non-specific binding by the secondary antibody, we omitted CSAb from the immunohistochemical protocol. We observed a total loss of signal, confirming that the staining was due to CSAb (Fig. S3). We also observed CSAb reactive deposits in non-neuronal cells in fALS subjects similar to previous reports (Fig. 3C, 3D)^29^. In SOD1-linked fALS patients, staining appeared across varied cellular compartments and cell types. Three of the four subjects showed cytoplasmic reactivity in the gray matter motor neurons and nuclear reactivity in non-neuronal cells. In non-SOD1 fALS subjects, similar cellular compartments in both neuronal and non-neuronal cells displayed CSAb reactive oligomers, further supporting the idea that SOD1 misfolding contributes to diverse ALS genotypes^30–37^. Surprisingly, in sALS subjects, CSAb reactivity was confined to motor neurons with no non-neuronal aggregates, indicating that the involvement of misfolded SOD1 may differ between sALS and fALS. This variation suggests that misfolded or aggregated SOD1 may exert its toxic effects through different cellular compartments and cell types, potentially reflecting diverse mechanisms of pathogenesis. Overall, the immunohistochemical analyses suggests that the SOD1 segment containing residues (28-38) is involved in ALS pathogenesis across different genotypes, possibly in the corkscrew conformation.

**Figure 3:**
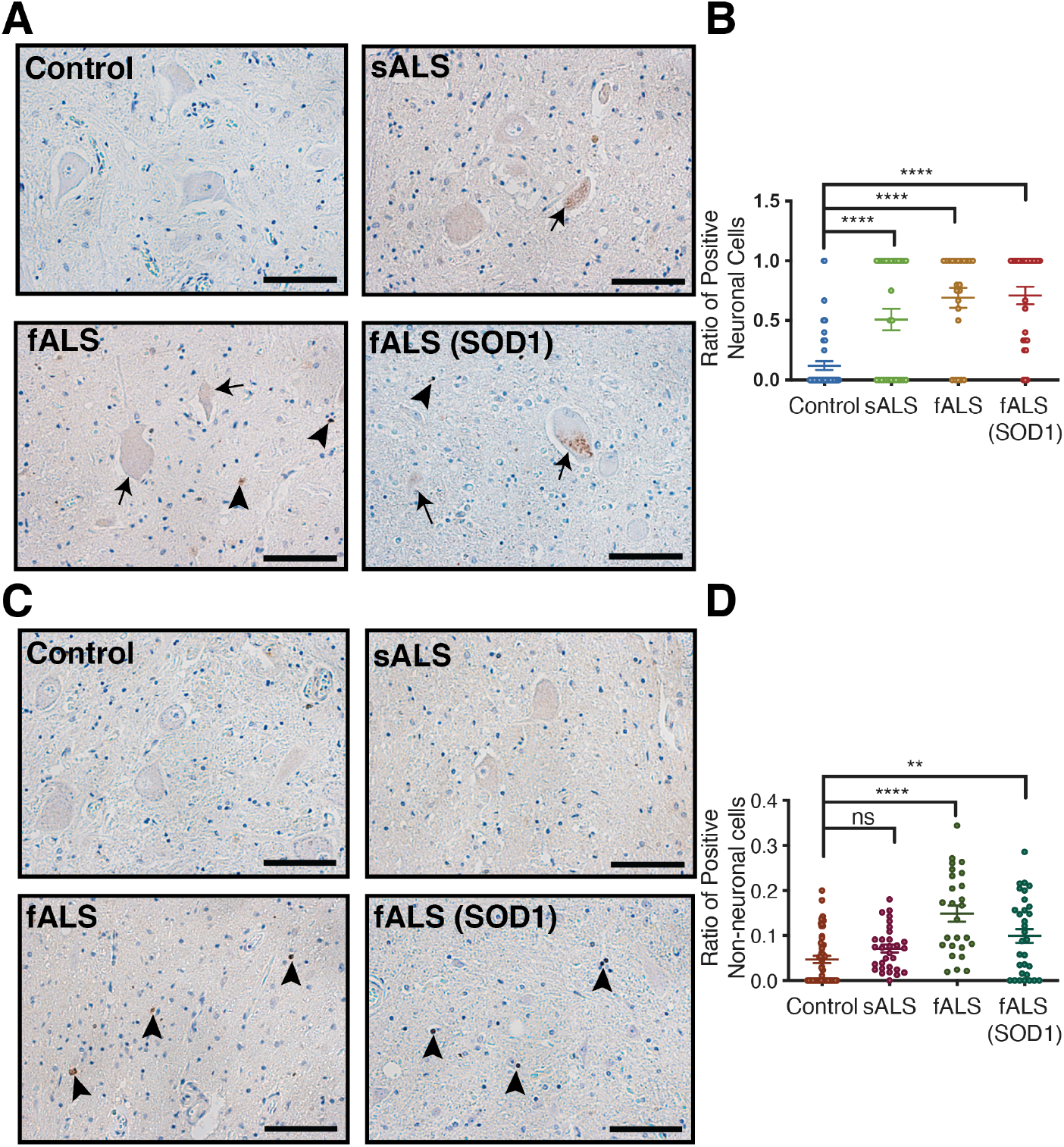
CSAb recognizes SOD1 oligomers in familial and sporadic ALS. **(A,C)** Immunohistochemical analyses of spinal cord sections of fALS, SOD1-linked fALS and sALS human subjects. Motor neurons of all ALS genotypes displayed CSAb reactive deposits while healthy controls displayed no reactivity. Intra-neuronal deposits were detected in all ALS genotypes ((sALS, fALS, SOD1-linker fALS)) while non-neuronal cells displayed deposits in fALS subjects but not sALS or healthy subjects. **(B,D)** Quantification of CSAb reactivity. Five fields of view were selected for each subject, and the ratio of cells displaying CSAb reactivity was quantified. In total, 27 subjects were analyzed. Results are shown as mean and standard deviation. Statistical significance was determined by ANOVA. ns not significant, **<.01, **** <.0001.

## Discussion

Our study demonstrates the critical role of corkscrew oligomers in ALS pathogenesis and highlights the utility of a monoclonal antibody, CSAb, designed to target this structure. We show that CSAb selectively binds to toxic oligomeric forms of SOD1, distinguishing them from native and fibrillar forms, and reduces their cytotoxicity in motor neuron cultures. Furthermore, CSAb detects misfolded SOD1 in both familial and sporadic ALS patient tissues, suggesting that the corkscrew epitope is a shared pathological feature across ALS subtypes. Our findings suggest CSAb can also be used as a probe for studying SOD1 pathology.

CSAb recognizes a conformational epitope rather than the linear sequence of segment, 28-38. Under non-denaturing conditions such as dot blot binding assay and native gel electrophoresis, CSAb selectively detects oligomers. Under the denaturing conditions such SDS-PAGE, it does not recognize the metal-bound forms of the protein, although it can bind to apo-forms of the protein. Given SOD1’s exceptional stability, it may retain partial secondary structure even in harsh denaturing conditions such as SDS-PAGE, which leads to differential binding properties. Current models of cytotoxic SOD1 oligomers suggest corkscrew-like oligomers or non-native trimers. Based on our native gel electrophoresis results, which show that CSAb recognizes oligomers from dimers to hexamers, we propose that cytotoxic SOD1 oligomers present an epitope comprised of residues 28-38.

Immunohistochemical analyses of patient-derived tissues suggest SOD1 aggregation plays a role in both inherited and non-inherited forms of ALS. This suggests that toxic SOD1 oligomers may arise through multiple mechanisms, including genetic mutations in fALS or post-translational modifications and cellular stress in sALS^38–42^. Furthermore, the detection of oligomeric SOD1 in non-SOD1 fALS cases suggests that SOD1 misfolding may occur as a downstream pathological event, in cases where SOD1 mutations are absent. This highlights a shared role for SOD1 toxicity across ALS genotypes and emphasizes the broader relevance of SOD1 misfolding in ALS.

The distinct cytoplasmic and nuclear staining patterns observed with CSAb underscores the complexity of SOD1 pathology in ALS. In SOD1-linked fALS cases, the presence of cytoplasmic aggregates within motor neurons aligns with the established role of SOD1 aggregates in motor neuron degeneration^43,44^. In contrast, the nuclear reactivity observed in non-neuronal cells, such as astrocytes and microglia, suggests a non-cell autonomous mechanism of toxicity^30–37^. Non-neuronal cells are increasingly implicated in ALS progression, with evidence showing that astrocytes expressing mutant SOD1 can secrete neurotoxic factors that exacerbate motor neuron degeneration^31,32,34^. Our findings suggest the misfolded SOD1 is prevalent in diverse ALS genotypes but may exert its toxic effects through different mechanisms.

CSAb could be used as a versatile tool in the study and treatment of ALS. As a probe, CSAb can detect toxic oligomeric SOD1 conformations in patient tissues, aiding in the development of diagnostic biomarkers for both familial and sporadic ALS. Its specificity for the corkscrew epitope allows it to distinguish pathogenic forms of SOD1 from native or fibrillar forms, a key advantage over existing antibodies. Additionally, CSAb’s ability to detect misfolded SOD1 in various cellular compartments and cell types suggests its utility in studying the distribution and dynamics of SOD1 across ALS subtypes. This may provide insights into the progression of SOD1-mediated cytotoxicity, including the interaction of the toxic species with other cellular components. CSAb could also be applied as a tool for isolating and characterizing oligomeric SOD1, enabling detailed structural studies. This could advance our understanding of the molecular mechanism of SOD1 aggregation. Finally, CSAb holds promise as a therapeutic candidate. Unlike current therapeutic strategies, such as antisense oligonucleotides (ASOs) that cause widespread SOD1 suppression, CSAb specifically targets pathogenic forms, offering a more precise approach that could reduce off-target effects and preserve native SOD1 function.

Oligomeric intermediates are increasingly recognized as toxic drivers in disorders such as Alzheimer’s, Parkinson’s, and Huntington’s disease. Our approach of identifying a minimal, structurally unique epitope within a misfolded protein and generating a highly selective antibody could be applied to these diseases as well.

## Materials and Methods

### Expression and purification of SOD1 constructs

All SOD1 constructs were expressed recombinantly using an E. coli system. All recombinant proteins are based on a SOD1 background strain that carries a C6A, C111S double mutation to simplify the purification. The SOD1 gene was inserted into the pET22b vector from Novagen with NcoI and SalI restriction enzyme digestion sites (gift from Professor Joan Valentine’s lab at UCLA). Mutations in the SOD1 gene were made using the QuickChange Site-Directed mutagenesis Kit (Stratagene). The plasmid was transformed into the BL21(DE3)Gold (Agilent Technologies) expression strain. For expression, 10 ml LB + Amp (50µg/mL) was inoculated from frozen stock and grown overnight. 10 ml of starting culture was added to a 2L flask of 1L LB + Amp (50μg/mL) and grown for 3 hours at 37 °C to OD600 = 0.6. IPTG was then added to 1 mM and Zn2+ was added to 0.05 mM to induce protein expression, which continued for an additional 3 hours. The bacterial pellet was collected by centrifugation at 4000 rpm for 10 mins.

To purify SOD1 from the bacteria, osmotic shock was used to release proteins including SOD1 from the periplasm. First, the cell pellet was resuspended in 30 mL of 30 mM Tris-HCl pH 8 with 20% sucrose and stirred slowly for 20 mins. The cells were collected by centrifugation at 10,000 × g at 4°C for 10 mins. The pellet was then resuspended in 30 mL of ice-cold water and stirred slowly for 20 mins on ice to release periplasmic proteins. Next, cell debris was removed by centrifugation at 4°C for 10 mins at 10,000 × g. The contaminant proteins were then removed by precipitation with ammonium sulfate. 0.326 g/ml (NH4)2SO4 was added to the supernatant, and the solution was stirred at 4°C for 45 mins. The protein was purified out of ammonium sulfate by separation over a phenyl sepharose column. The column was equilibrated with buffer A (2 M (NH4)2SO4, 0.15 M NaCl, 0.05 M KH2PO4, pH 7.0, adjusted with 1 M KOH) and buffer B (0.15 M NaCl, 0.05 M KH2PO4, pH 7.0, adjusted with 1 M KOH). The protein eluted at 6-30% buffer B. The protein was then concentrated and further purified by size exclusion chromatography on a silica G3000 column (Tosoh Bioscience). The column buffer comprised 0.1 M sodium sulfate, 25 mM sodium phosphate, and 1 mM sodium azide, pH 6.5.

Additional methods are provided in supplementary information.

## Acknowledgments

We thank members of the D.S.E. and B.N. laboratories for helpful discussions and Anand Panchal at Silverlake Research for antibody generation.

## Funding

This research was supported by HHMI and NIH 4R01AG029430. SS acknowledges support from the dissertation year fellowship.

## Author contributions

DM collected data, NK collected and analyzed data, BN analyzed data, MG analyzed data, DSE designed the experiments and analyzed data. HR analyzed data and wrote the manuscript. SS designed the experiments, collected and analyzed data and wrote the manuscript. All authors edited the manuscript.

## Notes

### Competing Interest Statement

The authors have declared no competing interest.

### Summary of Updates

Author Affiliation Updated For Mark Geisberg.

