## Supplementary Information for "A structure-guided antibody detects SOD1 oligomers in diverse ALS genotypes" (from another version of this preprint: version 1, 2025-05-08)

**Figures S1-S3**

**Table S1**

**Materials and Methods**

Figure S1

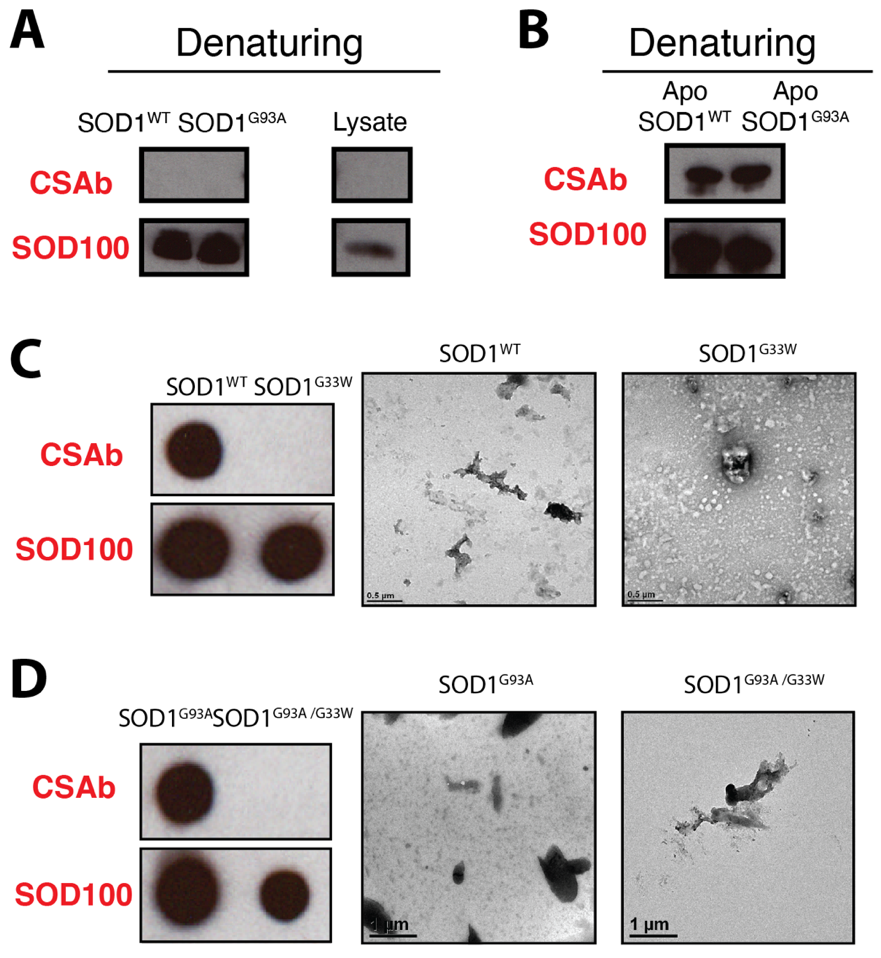

**Figure S1: CSAb recognizes the 28-38 epitope in apo-SOD1. (A)** Western blots of purified SOD1^WT^ and SOD1^G93A^ proteins and cell lysates show that CSAb does not recognize denatured protein. **(B)** Western blots of apo-proteins, SOD1^WT^ and SOD1^G93A^ showing CSAb reactivity. SOD100 was used to compare loading. **(C,D)** Dot blot binding assay of recombinantly purified SOD1 proteins (SOD1^WT^, SOD1G33W, familial mutant SOD1^G93A^ and SOD1^G93A/G33W^) with CSAb. Purified proteins were de-metalated and aggregated for 2 hours to form oligomers. CSAb recognizes SOD1^WT^ and SOD1^G93A^ mutant proteins but not SOD1^G33W^ mutants, suggesting that the integrity of the sequence is essential for CSAb reactivity.

Figure S2

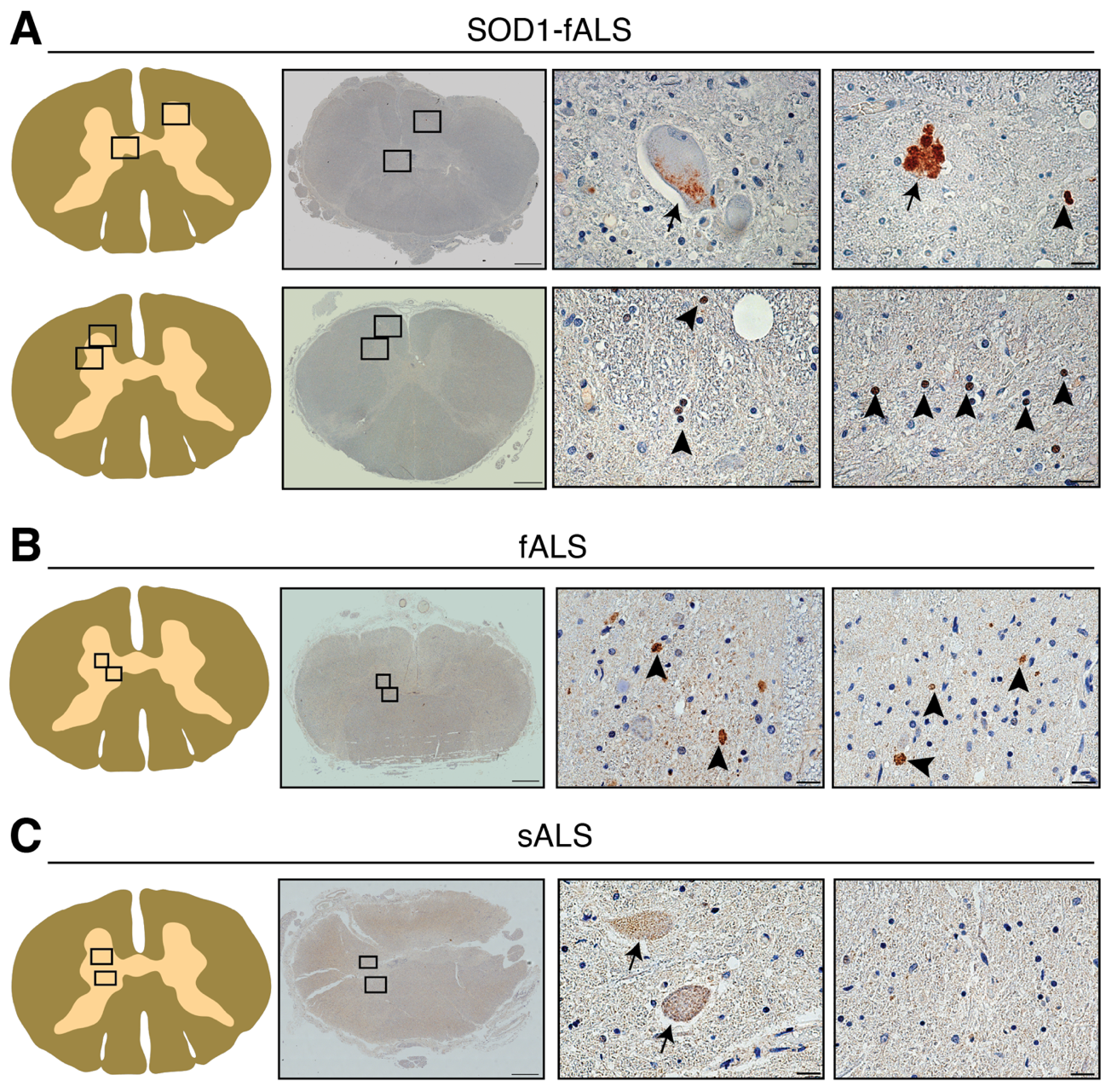

**Figure S2: CSAb reactivity in different ALS subjects.** Immunohistochemical micrographs of human ALS tissues probed for CSAb reactivity. **(A)** In two SOD1-fALS subjects, we found different reactivity patterns. In one subject (top row), we found predominantly cytoplasmic reactivity in the gray matter. In another subject (bottom row), we observed predominantly nuclear reactivity in non-neuronal cells in the gray matter. **(B)** In non-SOD1 linked fALS subjects, we observed both neuronal and non-neuronal deposits. **(C)** In sALS subject, we observed reactivity in motor neurons but not in the non-neuronal cells. Scale bar of entire tissue sections (in A, B, C) 1000 μm and 20 μm of higher magnifications.

Figure S3

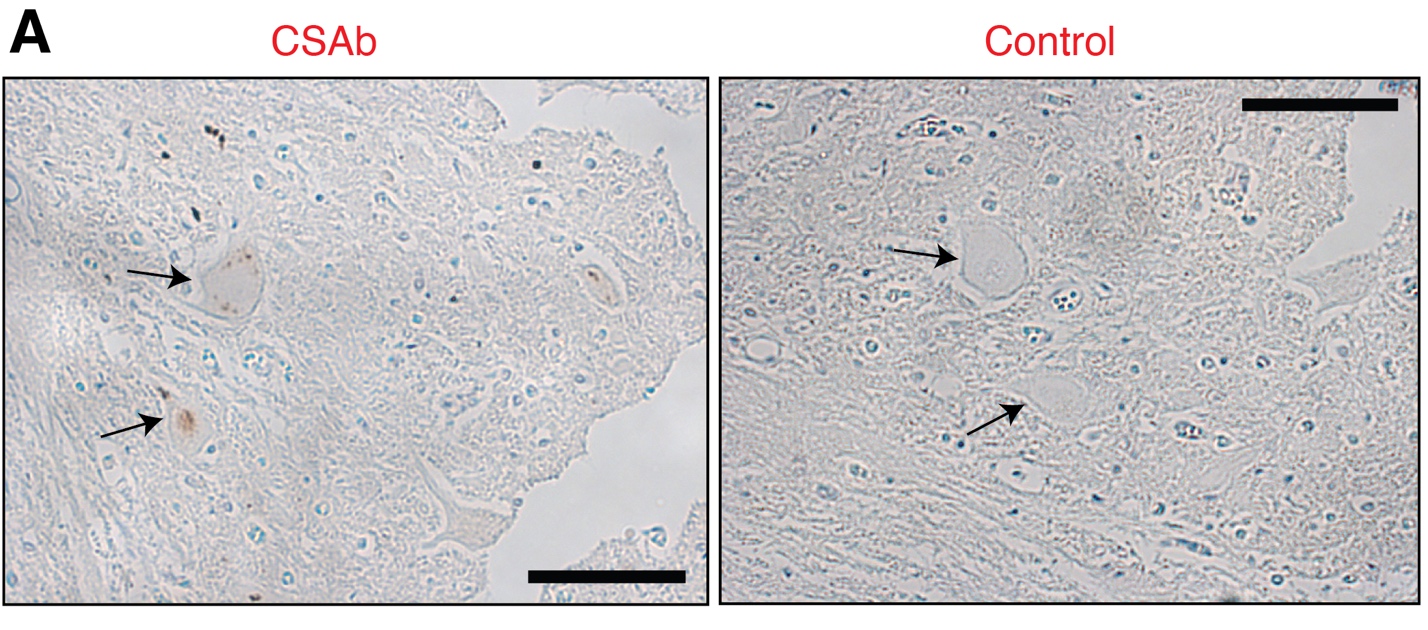

**Figure S3:** The specificity of CSAb was tested by removing the antibody from the immunostaining protocol. Notice the loss of intra-neuronal staining in the control sample, confirming that the staining is due to CSAb reactivity. Scale bar 20 μm.

**Table 1. Clinical information of samples used for CSAb reactivity.**

(NK – not known, NA – not applicable)

|  | **Clinical**  **Diagnosis** | **CSAb**  **(Non-neuronal)** | **CSAb**  **(Neuronal)** | **Pathologic diagnosis** | **Braak NFT stage** | **Thal amyloid phase** | **TDP-43** | **FHx** | **Age** | **Sex** | **Race** |
| --- | --- | --- | --- | --- | --- | --- | --- | --- | --- | --- | --- |
| 1 | ALS | 0.09 | 1 | ALS | 0 | 0 | Yes | No | 46 | M | Caucasian |
| 2 | ALS | 0.08 | 0 | ALS | 0 | 0 | Yes | No | 59 | F | Caucasian |
| 3 | ALS | 0.10 | 0.2 | ALS | 0 | 0 | Yes | No | 48 | M | Caucasian |
| 4 | ALS | 0.046 | 0.94 | ALS | I | 0 | Yes | No | 58 | F | Caucasian |
| 5 | ALS | 0.06 | 1 | ALS | I | 0 | Yes | No | 56 | F | Caucasian |
| 6 | ALS | 0.05 | 0 | ALS | NK | NK | NK | NK | 36 | M | Not specified |
| 7 | Control | 0.04 | 0.07 | Control | 0 | 0 | NA | No | 57 | M | Caucasian |
| 8 | Control | 0.09 | 0.50 | Control | 0 | 0 | No | No | 67 | M | Caucasian |
| 9 | Control | 0.09 | 0.27 | Control | III | 0 | NA | No | 71 | F | Caucasian |
| 10 | Control | 0.04 | 0 | Control | III | 1 | NA | No | 72 | M | Caucasian |
| 11 | Control | 0 | 0 | Control | NK | NK | NA | NA | 62 | F | Caucasian |
| 12 | Control | 0 | 0 | Control | NK | NK | NA | NA | 69 | F | Not specified |
| 13 | Control | 0 | 0 | Control | NK | NK | NA | NA | 72 | M | Not Specified |
| 14 | Control | 0.07 | 0.13 | Control | NK | NK | NA | NA | 57 | M | Not Specified |
| 15 | Control | 0.10 | 0.20 | Control | 0 | 0 | NA | NA | 92 | F | Not Specified |
| 16 | Control | 0.02 | 0 | Control | 0 | 0 | NA | NA | 80 | M | Not Specified |
| 17 | ALS | 0.20 | 0.46 | ALS, familial (SOD1 A4V) | I | 0 | No | Yes | 55 | F | Caucasian |
| 18 | ALS | 0.11 | 0.93 | ALS, familial (SOD1 G141X) | 0 | 0 | No | Yes | 42 | F | Caucasian |
| 19 | ALS | 0.14 | 0.87 | ALS, familial (SOD1) | 0 | 0 | No | Yes | 55 | M | Caucasian |
| 20 | ALS | 0.01 | 1 | ALS, familial (SOD1 A4V) | NK | NK | NK | NK | 68 | M | Not specified |
| 21 | ALS | 0 | 0.06 | ALS, familial (SOD1 A4V) | NK | NK | NK | NK | 65 | F | Not specified |
| 22 | ALS | 0.10 | 0.87 | ALS, familial (SOD1 A4V) | NK | NK | NK | NK | 21 | F | Not specified |
| 23 | ALS | 0.05 | 0.83 | ALS, familial (Chr9) | I | 0 | Yes | Yes | 58 | M | Caucasian |
| 24 | ALS | 0.21 | 0.74 | ALS, familial (Chr9) | I | 0 | Yes | No | 51 | F | Caucasian |
| 25 | ALS | 0.26 | 0.94 | ALS, familial (Chr9) | I | 0 | Yes | No | 49 | F | Caucasian |
| 26 | ALS-MCI | 0.09 | 0 | ALS, familial (Chr9) | I | 0 | Yes | No | 53 | M | Caucasian |
| 27 | ALS | 0.12 | 1 | ALS, familial (Chr9) | II | 0 | Yes | Yes | 58 | F | Caucasian |

**Materials and Methods**

**Expression and purification of SOD1 constructs**

All SOD1 constructs were expressed recombinantly using an E. coli system. All recombinant proteins are based on a SOD1 background strain that carries a C6A, C111S double mutation to simplify the purification. The SOD1 gene was inserted into the pET22b vector from Novagen with NcoI and SalI restriction enzyme digestion sites (gift from Professor Joan Valentine’s lab at UCLA). Mutations in the SOD1 gene were made using the QuickChange Site-Directed mutagenesis Kit (Stratagene). The plasmid was transformed into the BL21(DE3)Gold (Agilent Technologies) expression strain. For expression, 10 ml LB + Amp (50µg/mL) was inoculated from frozen stock and grown overnight. 10 ml of starting culture was added to a 2L flask of 1L LB + Amp (50μg/mL) and grown for 3 hours at 37 °C to OD600 = 0.6. IPTG was then added to 1 mM and Zn2+ was added to 0.05 mM to induce protein expression, which continued for an additional 3 hours. The bacterial pellet was collected by centrifugation at 4000 rpm for 10 mins.

To purify SOD1 from the bacteria, osmotic shock was used to release proteins including SOD1 from the periplasm. First, the cell pellet was resuspended in 30 mL of 30 mM Tris-HCl pH 8 with 20% sucrose and stirred slowly for 20 mins. The cells were collected by centrifugation at 10,000 × g at 4°C for 10 mins. The pellet was then resuspended in 30 mL of ice-cold water and stirred slowly for 20 mins on ice to release periplasmic proteins. Next, cell debris was removed by centrifugation at 4°C for 10 mins at 10,000 × g. The contaminant proteins were then removed by precipitation with ammonium sulfate. 0.326 g/ml (NH4)2SO4 was added to the supernatant, and the solution was stirred at 4°C for 45 mins. The protein was purified out of ammonium sulfate by separation over a phenyl sepharose column. The column was equilibrated with buffer A (2 M (NH4)2SO4, 0.15 M NaCl, 0.05 M KH2PO4, pH 7.0, adjusted with 1 M KOH) and buffer B (0.15 M NaCl, 0.05 M KH2PO4, pH 7.0, adjusted with 1 M KOH). The protein eluted at 6-30% buffer B. The protein was then concentrated and further purified by size exclusion chromatography on a silica G3000 column (Tosoh Bioscience). The column buffer comprised 0.1 M sodium sulfate, 25 mM sodium phosphate, and 1 mM sodium azide, pH 6.5.

**SOD1 demetallation (required for aggregation)**

The protein constructs purified by size exclusion chromatography were dialyzed in four buffers at 4°C to remove the bound metals. Protein preparations were put in a 10 kDa dialysis cassette and exchanged against buffer 1 twice (100mM Na Acetate pH 3.8, 10mM EDTA pH 8.0), followed by dialysis in buffer 2 (100mM Na Acetate pH 3.8, 100mM NaCl), buffer 3 (100mM Na Acetate pH 5.5, 100mM NaCl) for 4 hours, and finally in buffer 4 (20mM K phosphate pH7.0) for 12 hours each. To avoid metal contamination, all stock solutions (except for EDTA) were treated with Chelex 100 resin (Bio-Rad) to remove trace metals and all beakers were rinsed with metal-free water.

**Protein and peptide aggregation**

Full-length SOD1: Following metal removal, proteins were concentrated with a 10kDa concentrator to 800 μM (monomer) stock concentration and stored at -80 °C in 50μL aliquots until further use. Aggregation assays were carried out in 1.5 mL microcentrifuge tubes (Fisher Scientific) under acidic conditions with 80 μM SOD1, 50 mM sodium acetate, 1 mM EDTA and 1 mM TCEP, pH 3.5. Preparations were agitated at 900 rpm at 37 °C for indicated times. These samples were then dialyzed with 10 mM potassium phosphate and 1 mM EDTA, pH 7.0 for 4 hours at 4 °C.

Peptide: Peptide segments were aggregated by incubation at 37°C for 4-12 hours with agitation in 50 mM tris-base buffer and added to cultured neurons.

**Differentiation of mESCs to motor neurons**

Hb9:eGFP mESCs were maintained and differentiated into motor neurons as previously described (13). Briefly, mESCs were first plated on gelatin to remove mouse embryonic fibroblasts (MEFs) prior to differentiation, and then plated in 60 mm bacterial petri dishes in core motor neuron medium (DMEM/F12, Neurobasal, 10% Knockout Serum Replacement, Pen-Strep, Glutamax) to induce embryoid body (EB) formation. Two days later, N2 supplement (1x), retinoic acid (1 μM; Sigma), and SAG (1 μM; Calbiochem) were added to the EBs. Media was changed every two days. After five days post-retinoic acid and SAG addition, Hb9:GFP EBs were dissociated using ice-cold 0.25% Trypsin-EDTA for 6 minutes at room temperature, followed by trituration in L-15 medium (Hyclone). Dissociated motor neurons were plated in core motor neuron medium with GDNF, BDNF, CNTF (all 10 ng/ml; Peprotech) on 96-well plates that were previously coated with poly-L-ornithine (0.01%; Sigma) and laminin (5 μg/ml; BD Biosciences) and incubated for 12-16 hrs before treatment with the SOD1 peptides or proteins. Approximately 80,000 cells were plated per well. All reagents are from Invitrogen, unless otherwise noted.

**Cytotoxicity Assays**

Unless stated otherwise, all full-length proteins were aggregated for 12 hours before adding to motor neurons at 8 μM final concentration. Motor neurons were incubated for 12-16 hrs and imaged (Axio Observer.D1 microscope, Zeiss), followed by a cell viability assay. Cell viability was measured using the commercially available MTT assay (Promega Cat#G4100) according to the manufacturer’s protocol with 4-hour incubation with the MTT reagent. All data were normalized using buffer as 100% viability and 0.2% SDS as 0% viability. Experiments were performed in triplicates and repeated a minimum of three times.

**Electron Microscopy**

3-5 μL sample was spotted directly on freshly glow-discharged carbon-coated electron microscopy grids (Ted Pella, Redding, CA). After 4 min incubation, grids were rinsed twice with 5μL distilled water and stained with 5μl of 2% uranyl acetate for 1 min. Excess uranyl acetate was removed by blotting and specimens were examined on a T-12 electron microscope at an accelerating voltage of 80 kV. Images were recorded digitally by wide angle (top mount) BioScan 600W 1 × 1K digital camera (Gatan, Pleasanton, CA).

**Dot Blot Assay**

3-5μL sample was spotted on a nitrocellulose membrane (Trans-Blot, Bio-Rad) followed by blocking with 10% fat-free milk in 1X TBST buffer (50 mM Tris, 150 mM NaCl, 0.05% Tween20) for 30 mins. Membranes were incubated with primary antibody - SOD100 (Enzo Life Sciences) at 1:1000 dilution, SODAb (gift by Prof. David Borchelt) at 1:5000 dilution, A11 (EMD Millipore) at 1:500, CSAb (in-house) at 1:500 diluted in 5% fat free milk in TBST buffer at room temperature for 1 hour. The membranes were washed three times in TBST buffer before incubating with HRP-linked secondary antibody - anti-mouse HRP secondary antibody (Life Technologies) at 1:10000 dilution, or anti-rabbit HRP secondary antibody (Life Technologies) at 1:10000 diluted in 5% fat-free milk in TBST buffer at room temperature for 1 hour. Pierce ECL Western Blotting Substrate, (Thermo Scientific, #32209), a chemiluminescent substrate was used for the detection of horseradish peroxidase (HRP) on the immunoblots following the manufacturer’s instructions. Detailed protocol can be found on <https://tools.thermofisher.com/content/sfs/manuals/MAN0011536_Pierce_ECL_West_Blot_Subs_UG.pdf>

**Antibody generation**

Mice were immunized with 10 μg of synthetic peptide KVKVWGSIKGL dissolved in PBS and incubated 12 hours at 4°C. Boosts were performed bi-weekly for 4-8 weeks with the same dosages. Mice were sacrificed and splenocytes were fused with Sp2/0 myeloma cells (American Type Culture Collection cell line CRL-1581, Manassas, VA) (21). Following HAT selection, surviving clones were cultured in microwell plates and selected for production of IgG binding to peptide KVKVWGSIKGL. These screens were performed by indirect ELISA on culture supernatants and confirmed by dot blots with KVKVWGSIKGL peptide bound to nitrocellulose. Monoclonal antibody (MAb) CSAb was identified by these criteria and the clone was stabilized by limiting dilution subcloning. MAb was produced from bulk hybridoma culture in medium with serum depleted of bovine IgG and purified by Protein G chromatography (Pierce Biotechnology, Rockford, IL).

**Blue native gel electrophoresis**

Samples were prepared by mixing with 4X Native PAGE sample buffer. For more details refer to the protocol for the Novex Native PAGE Bis-Tris Gel System (Thermo Fisher) (https://tools.thermofisher.com/content/sfs/manuals/nativepage_man.pdf). Briefly, equal amounts of proteins were loaded on 3-12% Bis-Tris native gels with light blue cathode buffer at 150 V for 2 hours. The protein was transferred to PVDF membranes using iBlot dry blotting system (Thermo Fisher) at 20V for 6 mins. Membranes were washed with methanol to remove Coommassie Blue and then washed with water. Membranes were then blocked with 10% fat-free milk in TBST for 1 hour at room temperature. Membranes were then washed three times with TBST and incubated with the primary antibody – SOD100 (Enzo Life Sciences) at 1:1000 dilution or CSAb (in-house) diluted 1:500 in 5% fat-free milk in TBST for 1 hour. Membranes were then washed with TBST three times and incubated with the secondary antibody – anti-mouse HRP secondary antibody (Life Technologies) at 1:10000 dilution or anti-rabbit HRP secondary antibody (Life Technologies) at 1:10000 dilution in 5% fat-free milk in TBST). Membranes were washed again three times with TBST and the films were developed following the protocol as described in the Kit (Thermo Scientific Pierce ECL Western Blotting Substrate, #32209).

**ELISA**

100 μL of protein solution (native, oligomers and fibrils prepared as described above) was added to each well of 96-well microplates (in triplicates) and incubated at 4ºC overnight without shaking. Blank wells (without any protein sample) were used as negative control. Wells were blocked by adding 200 μL of SuperBlock T20 blocking buffer and incubated at room temperature (RT) with shaking for 30 minutes. Wells were washed three times by incubating with 100 μL of TBST for 5 minutes at RT. CSAb or SOD100 were added at 1:500 dilution in TBST and incubated at RT for 1 hour with shaking. The wells were washed three times by incubating with 100 μL of TBST for 5 minutes at RT. Anti-mouse HRP secondary antibody (Life Technologies) at 1:10000 dilution or anti-rabbit HRP secondary antibody (Life Technologies) at 1:10000 dilution were added and incubated at RT for 1 hour with shaking. Wells were washed three times with TBST. 100 μL of TMB-ELISA solution (Thermo Scientific) was added and incubated for 5 minutes at RT. TMB substrate was neutralized with 100 μL 2M H_2_SO_4_. Absorbance was then measured at 450 nm.

**Immunohistochemistry**

Slides were deparaffinized by xylene followed by a series of washes in ethanol. After a wash under tap water, the slides were incubated in 3% hydrogen peroxide in methanol for 10 minutes. The slides were washed with distilled water and incubated for 10 minutes in EDTA Solution pH 8 (Invitrogen, #005501) at 95°C using a vegetable steamer. Following cool down to room temperature and washing with PBST (phosphate buffered saline containing 0.05% Tween-20), the slides were incubated at room temperature for 2 hours with anti-mouse CSAb antibody diluted 1:100 followed by Post Primary reagent of the Bond Refine Polymer Detection Kit (Leica, DS9800), for 30 minutes. Slides were rinsed with PBST and incubated with the Polymer reagent of the Bond Refine Polymer Detection Kit (Leica, DS9800), for 30 minutes, followed by incubation with DAB (3,3'-Diaminobenzidine) for visualization. Slides were counterstained with Harris’ Hematoxylin, dehydrated in ethanol, and mounted with media. Five fields of view were imaged for each patient sample. Neuronal and non-neuronal cells were counted and presented as ratio of total cells counted. All analyses were done blind.

**Statistical Significance**

All statistical analyses were done using GraphPad Prism 7.0.

**Antibodies used**

The following antibodies were used in this study. SOD100 (EnzoLife sciences) at 1:1000 dilution, A11 (EMD Millipore) at 1:500 dilution, anti β-actin (Abcam) at 1:1000, CSAb (in-house) at 1:500 dilution, anti-mouse HRP secondary antibody (Life technologies) 1:10000 dilution, anti-rabbit HRP secondary antibody (Life technologies) 1:10000 dilution.
